## Extended Data Figure 1-9 for "Single-cell RNA sequencing across isogenic *FUS* and *TARDBP* ALS lines reveals a shared early mitochondrial dysfunction unique to motor neurons"

Schweingruber et al., 2025

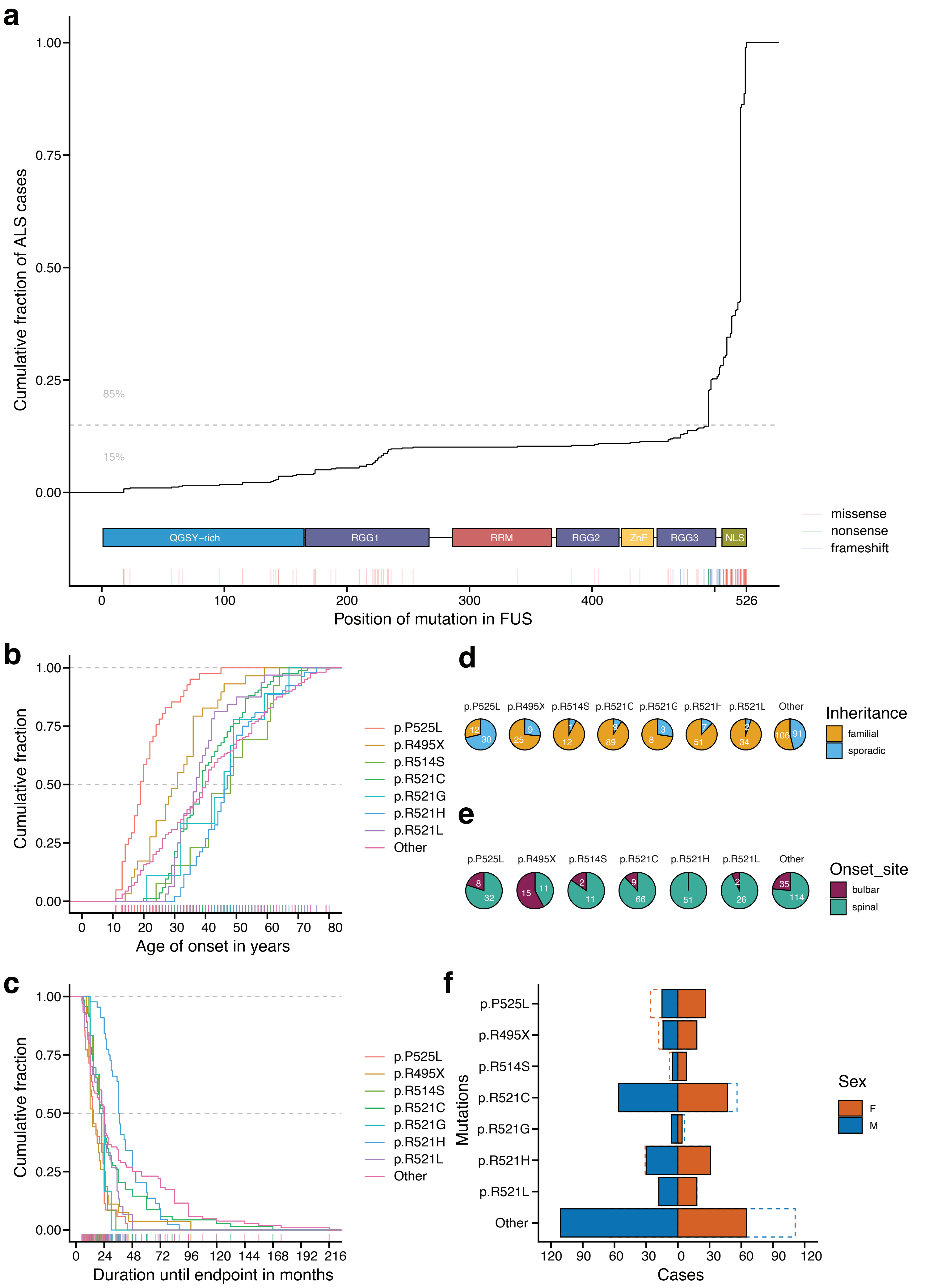

**Extended Data Figure 1. Meta-analysis of previously reported ALS cases with FUS mutations.** Clinical features from 502 individual cases with *FUS*-ALS were collected and aggregated from 115 articles or case reports published between 2009 and 2022. **a**, The ALS cases with FUS mutations were plotted as empirical cumulative density over the FUS protein. The C-terminal mutations account for approximately five sixth of the mutations reported in the dataset. **b-f**, Clinical features for *FUS* mutations with more than 10 cases in the dataset. The group 'Other' contains all the remaining mutations in the dataset. **b**, Age at onset in years is significantly different between FUS mutations (log-rank test, $\chi^2 = 175$, df = 7, $p < 2e\text{-}16$). **c**, Duration to endpoint (defined as respiratory failure or death) in months is significantly different between FUS mutations (log-rank test, $\chi^2 = 69.4$, df = 7, $p < 2e\text{-}12$). **d**, Inheritance of ALS (as familial or sporadic). **e**, The site of onset. **f**, Sex disparity.[1–115]

#### Extended Data Figure 2 Schweingruber et al., 2025

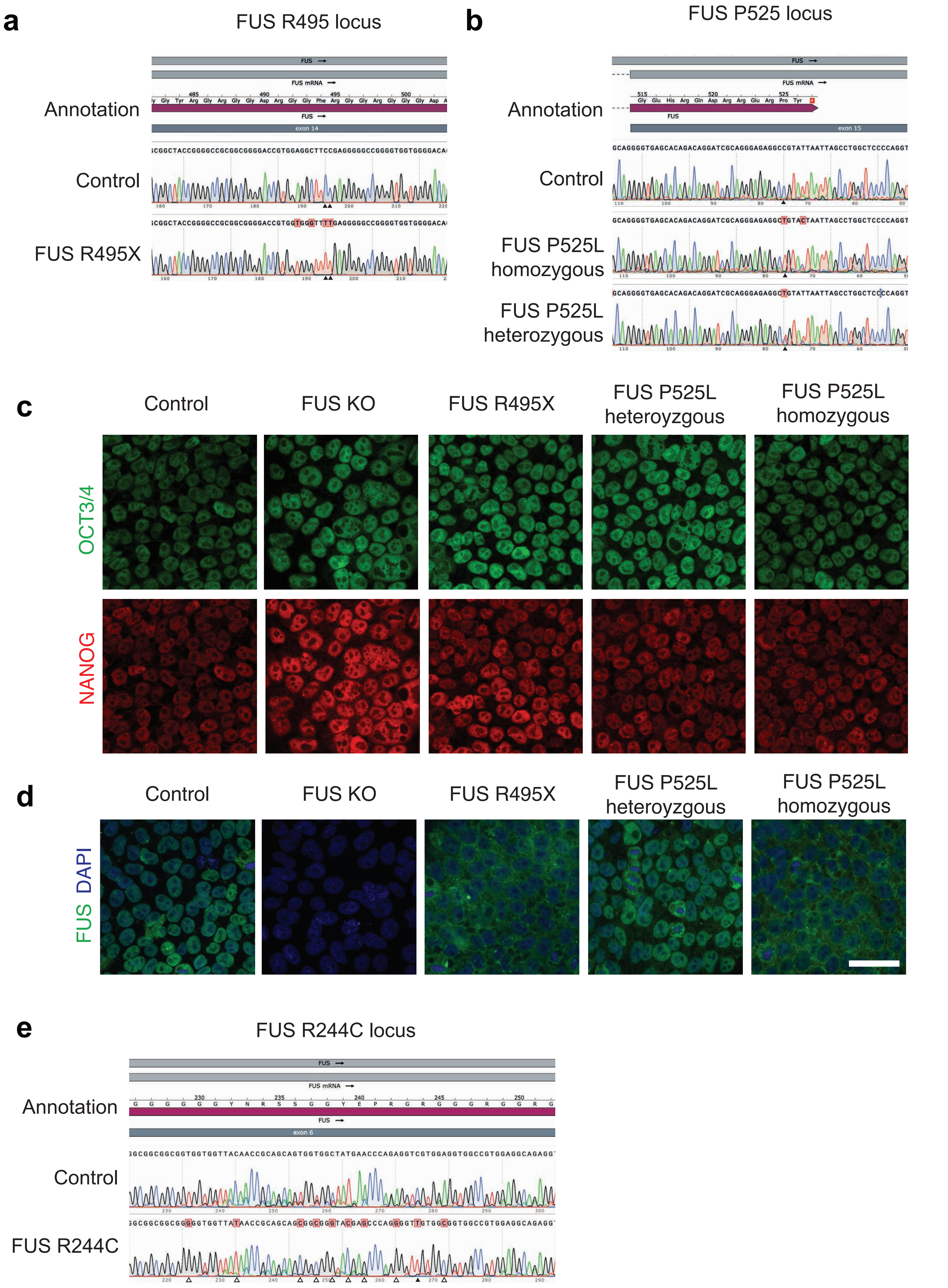

**Extended Data Figure 2. Characterization of human iPSC lines with FUS mutations.** Following genome editing of the DF6-9-9T.B control line, the editing sites were interrogated by dye terminator sequencing. **a**, Chromograms show the successful introduction of the R495X mutation in exon 14 of *FUS* in contrast to control cells. **b**, Chromograms show the introduction of the P525L mutation in exon 15 of *FUS* in a heterozygous and homozygous line. The editing of the FUS KO line was described previously [116,117]. **c**, Pluripotency of the genome-edited human iPSC lines was assessed with immunostainings with the stem cell markers OCT3/4 and NANOG. **d**, FUS expression and localization in the human iPSC lines was assessed by immunofluorescence with an antibody against N-terminal FUS and nuclear counterstaining with DAPI. **e**, Chromograms show the introduction of the R244C mutation in exon 6 of *FUS*. The white arrows indicate silent point mutations preventing re-editing of the locus whereas the black arrow indicates the target point mutation.

### Extended Data Figure 3

Schweingruber et al., 2025

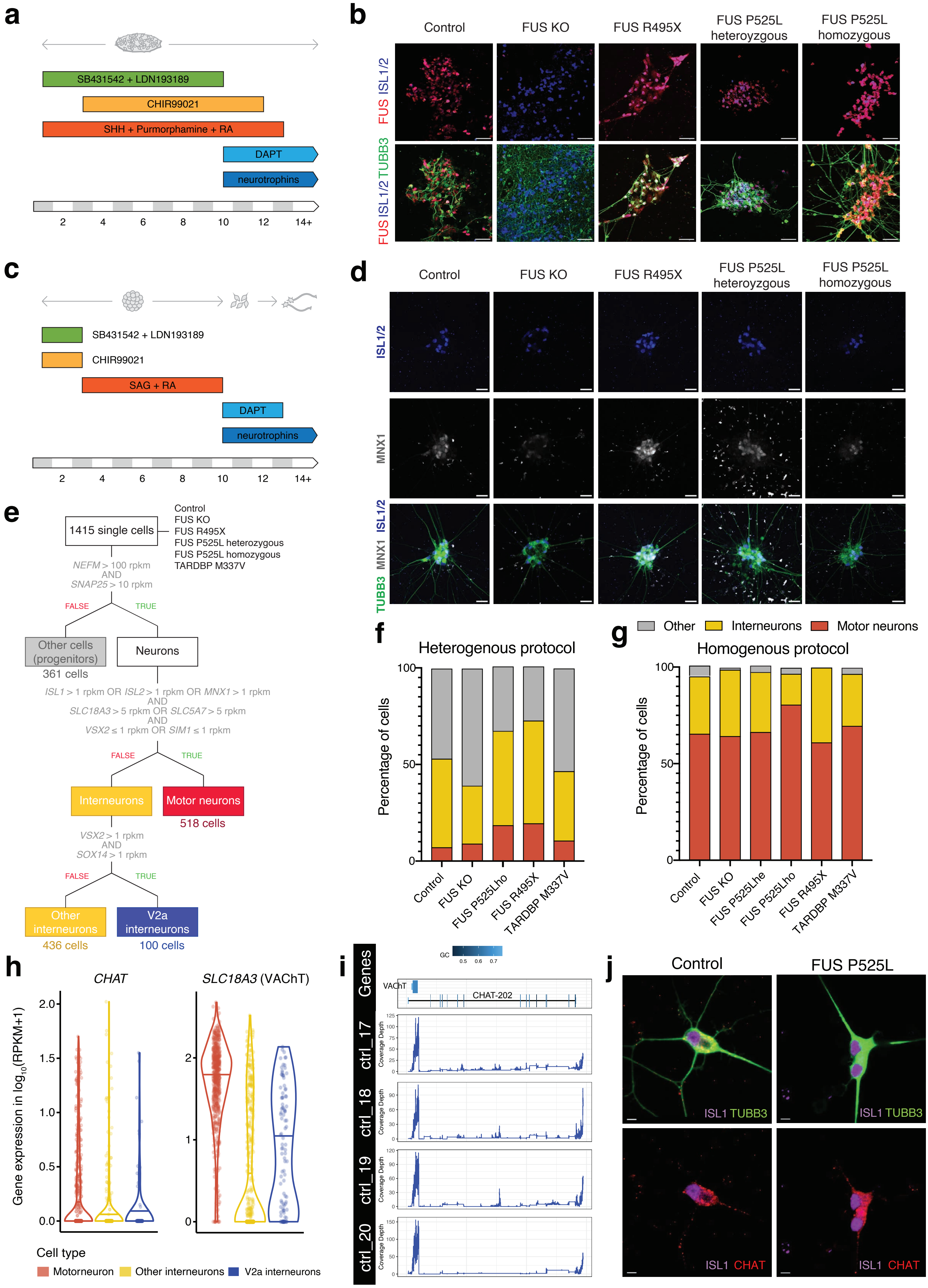

**Extended Data Figure 3. Differentiation of human iPSC to diverse spinal neurons. a**, Schematic of the differentiation protocol that yields a heterogenous population of spinal neurons including motor neurons. **b**, Expression of the motor neuron markers ISL1/2 and TUBB3 and FUS expression in the heterogenous protocol was assessed by confocal immunofluorescence microscopy. The scale bars measure 25 µm. **c**, Schematic of the differentiation protocol that yield a homogenous population of mostly spinal motor neurons. **d**, Expression of the motor neuron markers ISL1/2, MNX1 (Hb9) and TUBB3 in the homogenous protocol was assessed by confocal immunofluorescence microscopy. The scalebars measure 25 µm. **e,** The decision tree used for the cell type classification in our single cell RNA-Seq analysis. The specific expression cut-offs at each branch point are depicted. **f, g**, Following single cell RNA-Seq and cell-type classification, the proportions of motor neurons and interneurons and other cells (mostly neural progenitors) were calculated for the heterogenous **e**, and homogenous **f**, protocol, respectively. **h**, Expression of cholinergic markers *CHAT* and *SLC18A3* (VAChT) across cell types. **i**, Gene coverage plots for the nested CHAT-SLC18A3 genetic locus in four individual cells. **j**, Expression of the cholinergic marker CHAT in motor neurons (ISL1$^{+}$, Tuj1$^{+}$) specified with the homogenous protocol from control iPSC.

**Extended Data Figure 4** Schweingruber et al., 2025

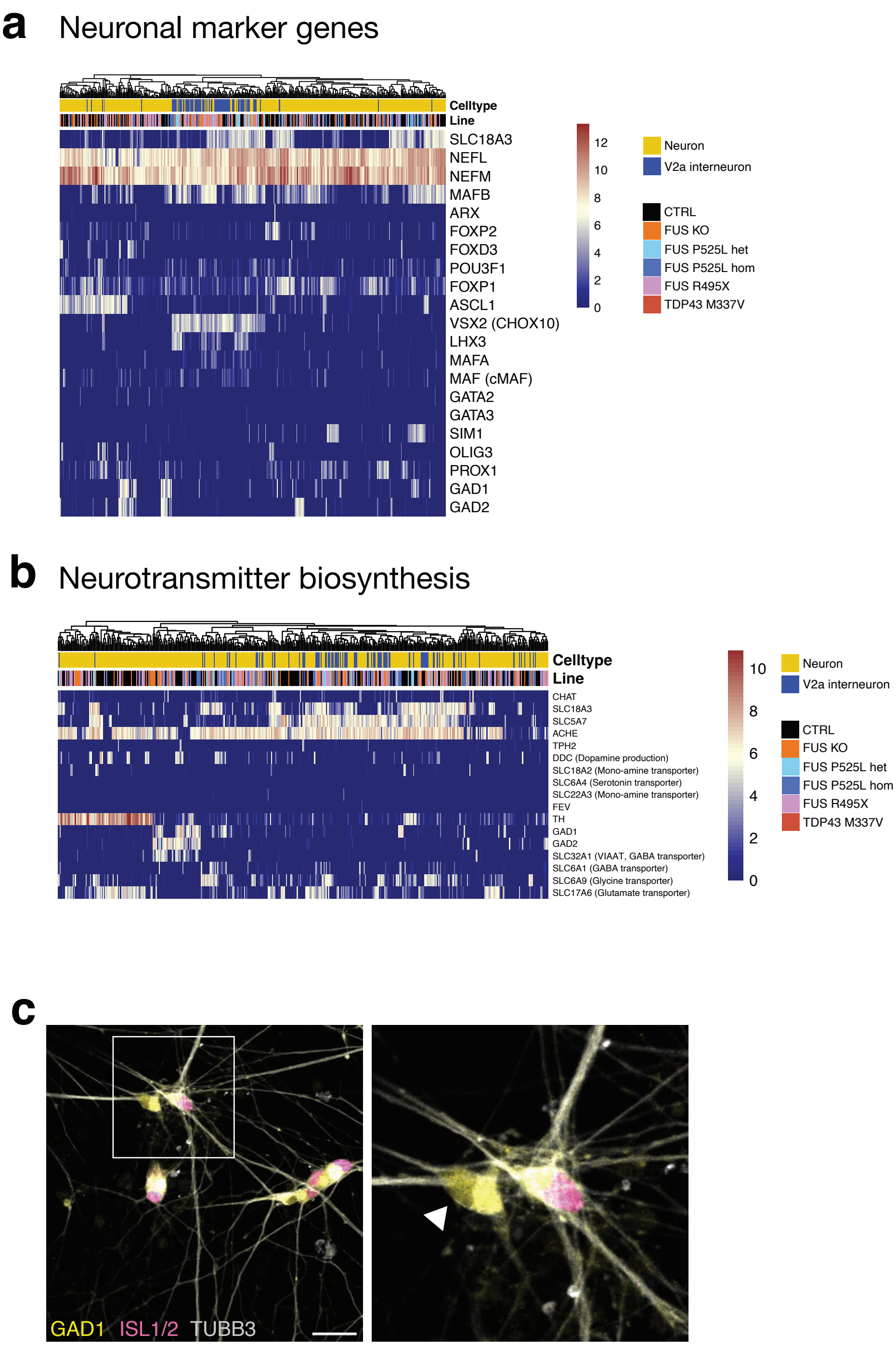

**Extended Data Figure 4. Several interneuron populations are present in the dataset.**

**a,** Heat map of the expression of selected markers in the other neurons and V2a interneurons. Hierarchical clustering groups the *VSX2*-expressing V2a interneurons. **b,** Heat map of neurotransmitter biogenesis and transporter gene expression in other neurons and V2a interneurons. Hierarchical clustering separates subgroups expressing *TH* (initiating the adrenergic and derivative neurotransmitter synthesis) and GABA-ergic neurons (*GAD1*, *GAD2*, *SLC32A1*, *SLC6A1*) among others. **c,** The presence of GABAergic interneurons (TUBB3+, ISLET1/2-, GAD1+, white arrow) among the motor neurons (TUBB3+, ISLET1/2+, GAD1+) was assessed by confocal immunofluorescence microscopy. The scale bars measure 30 µm.

### Extended Data Figure 5



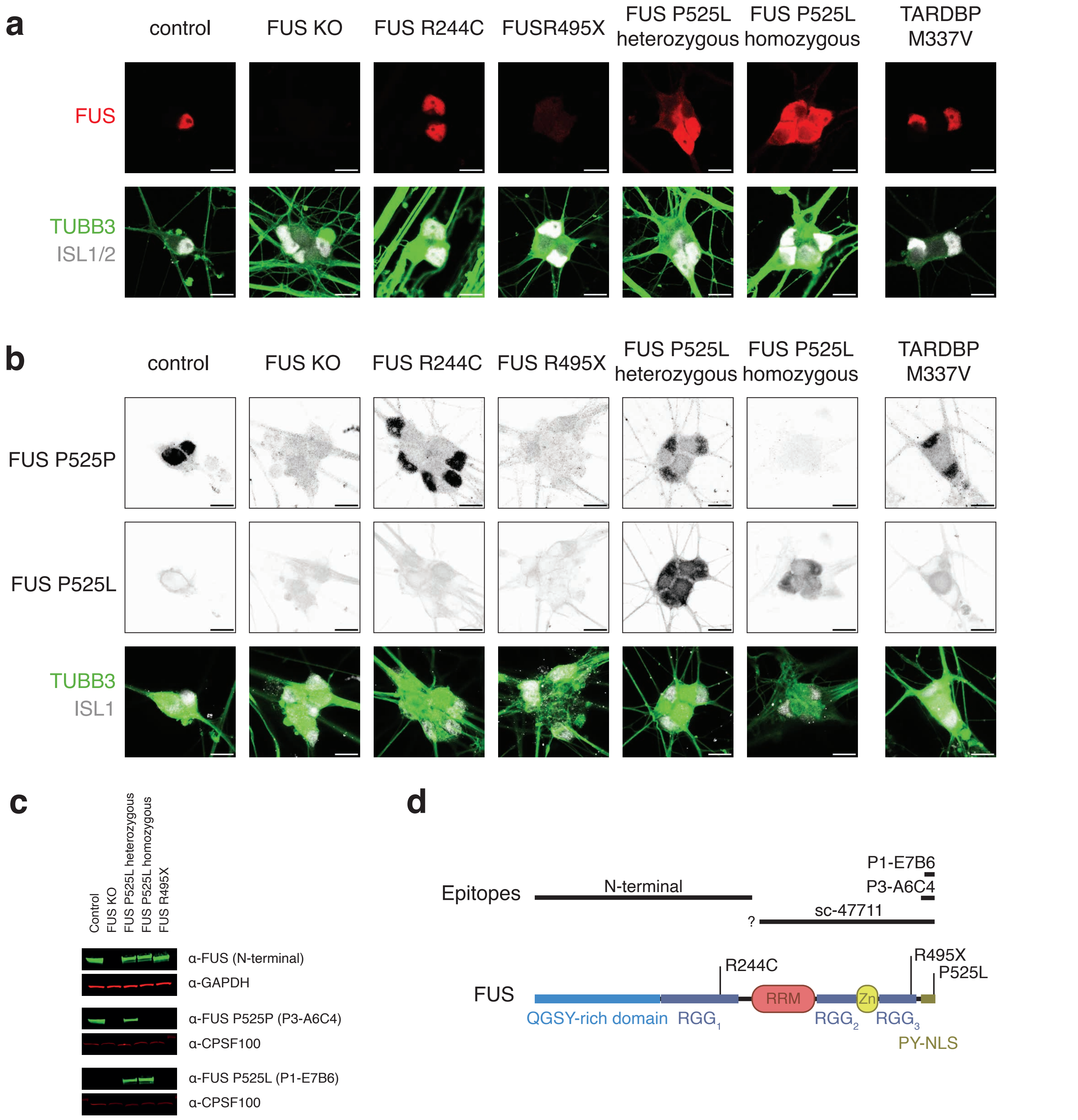

**Extended Data Figure 5. The intracellular localization of FUS is disturbed in ALS motor neurons.** **a**, Confocal fluorescence microscopy with immunostaining of C-terminal FUS in motor neurons (TUBB3+, ISL1/2+) at day 28 reveals cytoplasmic accumulation of FUS P525L and ablation in the FUS KO line. Scale bars are 10 µm. **b**, Interrogation with antibodies specific to FUS P525 epitopes allow us to separate the contribution of wild-type (P525P) and mutant (P525L) protein to the mis-localized pool of FUS in motor neurons at day 28. Scale bars are 10 µm. **c**, The specificity of the antibodies against the P525 epitope was assessed by western blot with lysates from the FUS mutant iPSC lines. **d**, Epitope map of the FUS antibodies used in this study including the FUS domain architecture as reference.

### Extended Data Figure 6

Schweingruber et al., 2025

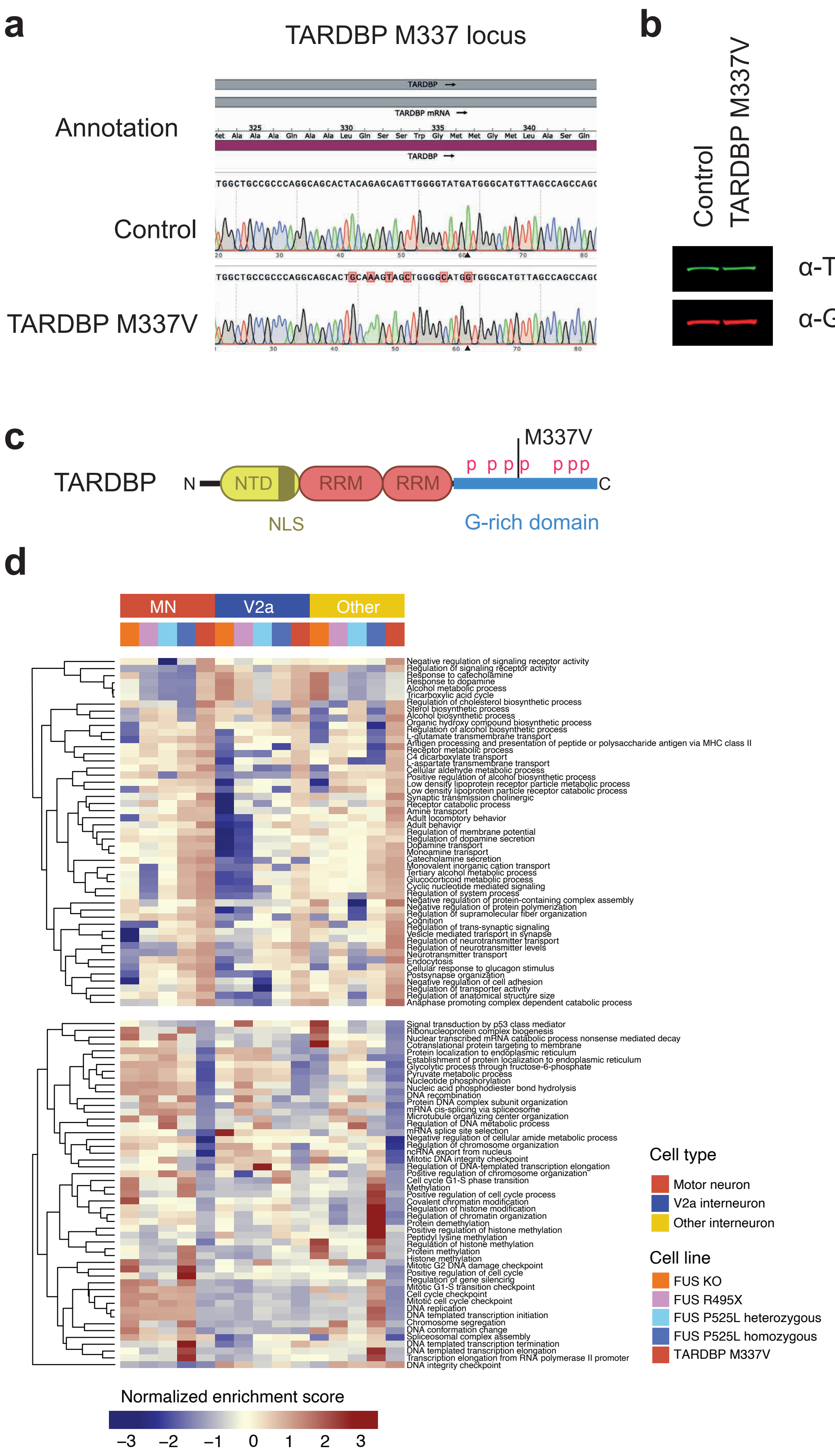

**Extended Data Figure 6. Characterization of the iPSC line with the TARDBP M337V mutation.** Following genome editing of the DF6-9-9T.B control line, the editing site was interrogated by dye terminator sequencing. **a,** Chromogram of the TARDBP M337 locus show the successful editing in the human iPSC line. **b,** Expression of TARDBP was assessed by western blot in lysates from control and mutant human iPSC lines. **c,** Domain architecture of TARDBP indicating the phosphorylation sites and location of the M337V mutation. **d**, Heatmaps display the normalized enrichment scores from GSEA of the top 25 up- and downregulated biological processes uniquely regulated in TARDBP M337V motor neurons. Motor neurons are compared to V2a or other interneurons to show specific or stronger regulation of these pathways in motor neurons.

### Extended Data Figure 7

Schweingruber et al., 2025

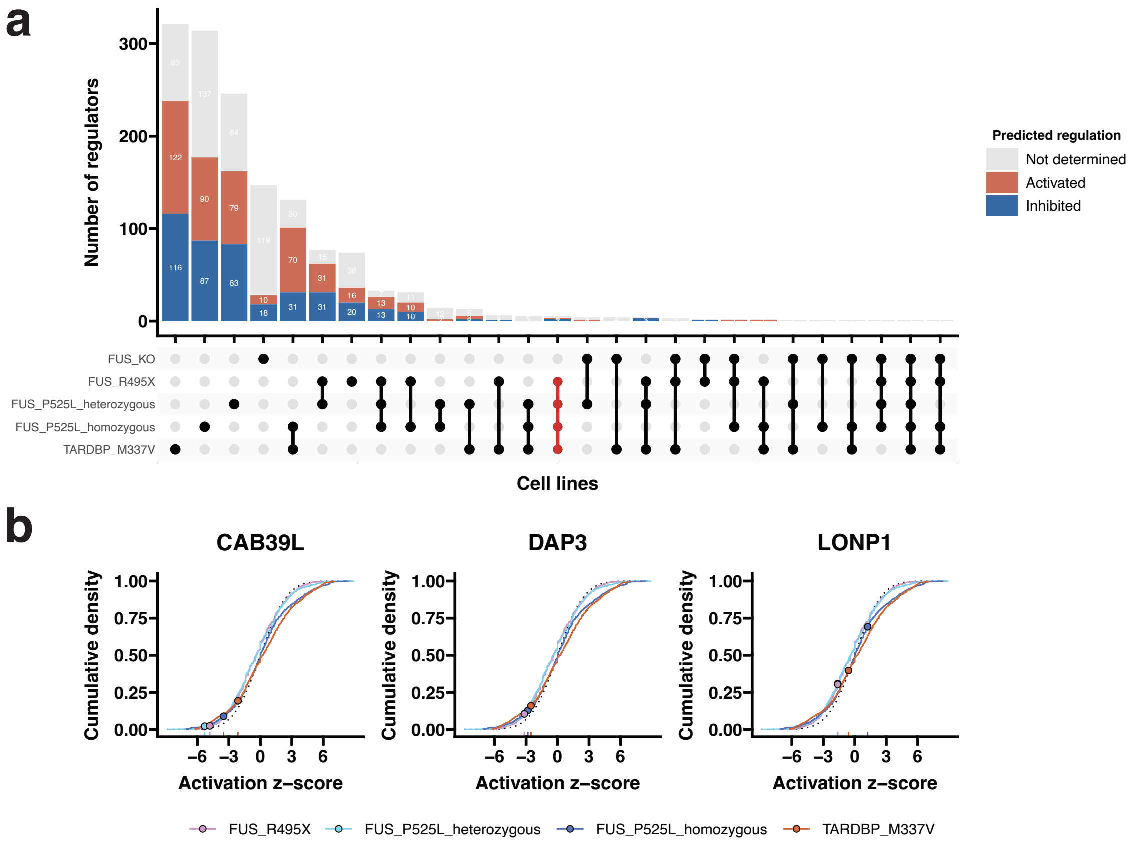

**Extended Data Figure 7. Upstream regulators of shared dysregulation in ALS lines.**

Upstream regulators that could explain the expression changes in the individual ALS-mutant motor neurons compared to the control were analyzed in the IPA tool set. **a,** The upset plot shows the intersection of upstream regulators with a network bias-corrected p-value < 0.001 separated according to their predicted regulation. The upstream regulator set that is shared across ALS lines is highlighted in red. **b,** The activation z-score for the protein-coding upstream regulators in the shared set are shown against the set of all upstream regulators in cumulative density plots. While a negative activation z-score indicates inhibition of the regulator, a positive score indicates its activation. CAB39L, DAP3, and LONP1 are all mitochondrial genes.

### Extended Data Figure 8

Schweingruber et al., 2025

**a**

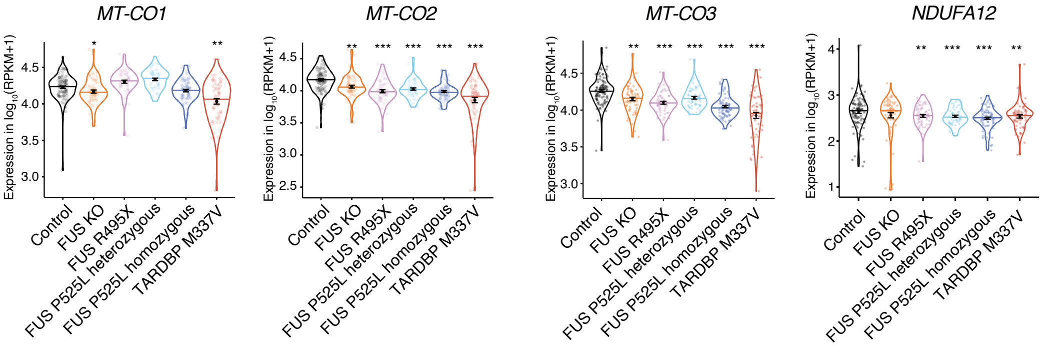

**Extended Data Figure 8. Expression of selected mitochondrial encoded genes.**

Violin plots showing normalized expression values (RPKM) of MT-CO1, MT-CO2, MT-CO3, and NDUFA12 in motor neurons generated with the homogeneous protocol. Error bars show mean ± SEM. The significance levels of the p-values are * $p < 0.05$, ** $p < 0.01$, *** $p < 0.001$ vs. control, derived from DESeq2 differential gene expression.

### Extended Data Figure 9

Schweingruber et al., 2025

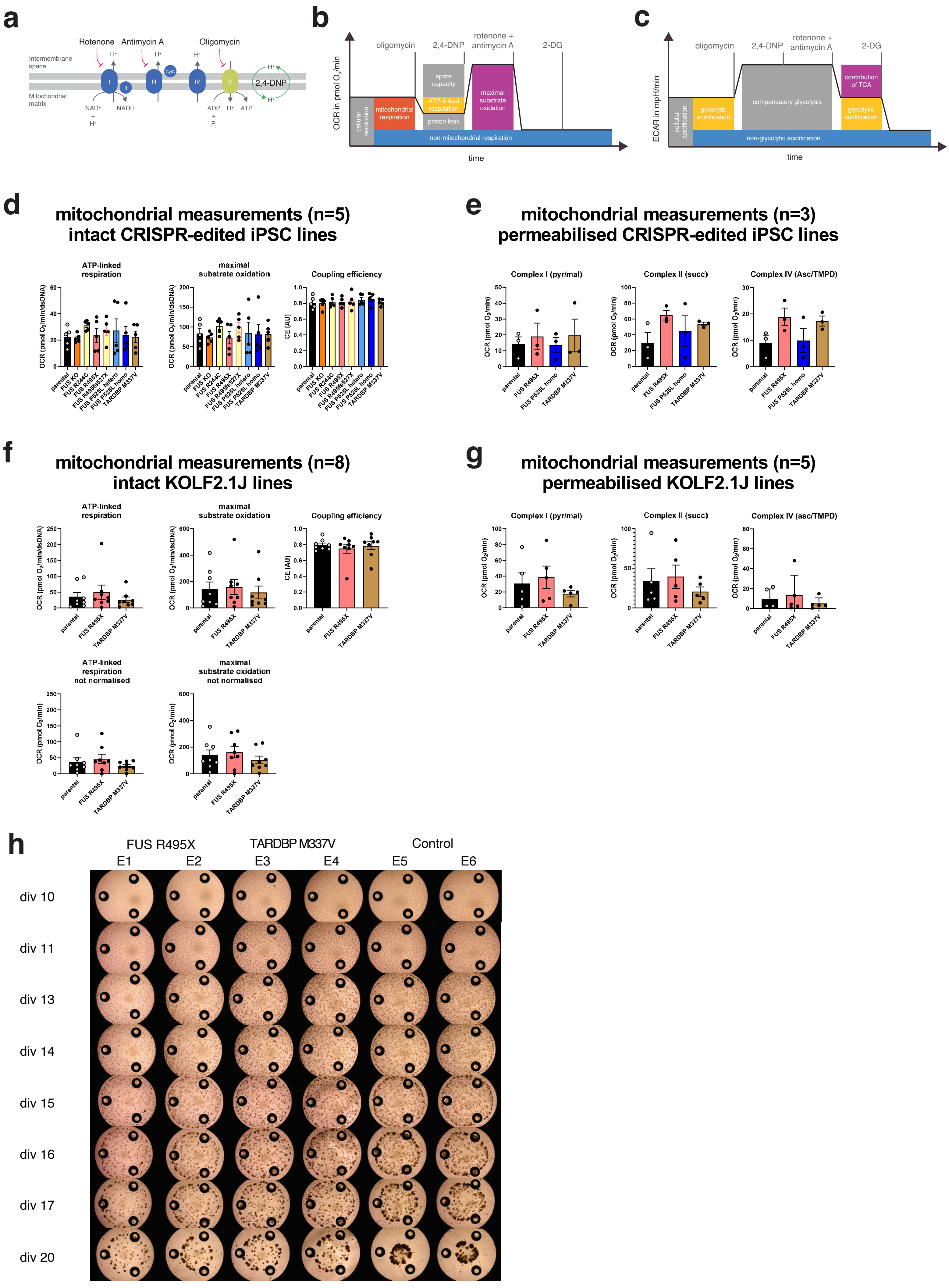

**Extended Data Figure 9. Bioenergetic profiling of mitochondria in motor neurons.** Bioenergetic profiling was conducted in motor neurons from the homogenous protocol at day 21. **a**, Specific mitochondrial inhibitors (oligo: oligomycin, R: rotenone, AA: antimycin A) and the uncoupler 2,4-Dinitrophenol (2,4-DNP), as depicted in the graph, and D-glucose metabolism inhibitor 2-deoxyglucose (2-DG), were used to analyzed oxygen consumption (OCR) and extracellular acidification (ECAR) using a XF96 extracellular flux analyzer as described in the Methods section. **b-c,** Partitioning of OCR (**b**) and ECAR (**c**) into different functional modules as depicted in the scheme (TCA: tricarboxylic acid cycle). **d-e,** Key metabolic parameters in intact and permabilized motor neurons derived from the DF-6-9-9T.B iPSC lines used in the single cell RNA sequencing. **f-g,** Key metabolic parameters in intact and permabilized motor neurons derived from KOLF2.1J iPSC lines. Data are the mean ± SEM of three to eight independent differentiations as indicated. * $p < 0.05$, + $p < 0.001$, # $p < 0.0001$ vs. control, One-way ANOVA (posthoc: Dunnett's). **h,** Representative images from motor neuron cultures in the KOLF2.1J background leading up to the bioenergetic profiling.

### Extended Data References


1. Chiò, A. *et al.* Two Italian kindreds with familial amyotrophic lateral sclerosis due to FUS mutation. *Neurobiol Aging* **30**, 1272–1275 (2009).
2. Kwiatkowski, T. J. *et al.* Mutations in the FUS/TLS Gene on Chromosome 16 Cause Familial Amyotrophic Lateral Sclerosis. *Science (1979)* **323**, 1205–1208 (2009).
3. Bäumer, D. *et al.* Juvenile ALS with basophilic inclusions is a FUS proteinopathy with FUS mutations. *Neurology* **75**, 611–8 (2010).
4. Blair, I. P. *et al.* FUS mutations in amyotrophic lateral sclerosis: clinical, pathological, neurophysiological and genetic analysis. *J Neurol Neurosurg Psychiatry* **81**, 639–45 (2010).
5. Bosco, D. A. *et al.* Mutant FUS proteins that cause amyotrophic lateral sclerosis incorporate into stress granules. *Hum Mol Genet* **19**, 4160–75 (2010).
6. Broustal, O. *et al.* FUS mutations in frontotemporal lobar degeneration with amyotrophic lateral sclerosis. *J Alzheimers Dis* **22**, 765–9 (2010).
7. Corrado, L. *et al.* Mutations of FUS gene in sporadic amyotrophic lateral sclerosis. *J Med Genet* **47**, 190–194 (2010).
8. Damme, P. van *et al.* The occurrence of mutations in FUS in a Belgian cohort of patients with familial ALS. *Eur J Neurol* **17**, 754–6 (2010).
9. DeJesus-Hernandez, M. *et al.* De novo truncating FUS gene mutation as a cause of sporadic amyotrophic lateral sclerosis. *Hum Mutat* **31**, E1377-89 (2010).
10. Groen, E. J. N. *et al.* FUS Mutations in Familial Amyotrophic Lateral Sclerosis in the Netherlands. *Arch Neurol* **67**, 224–30 (2010).
11. Hewitt, C. *et al.* Novel FUS/TLS mutations and pathology in familial and sporadic amyotrophic lateral sclerosis. *Arch Neurol* **67**, 455–61 (2010).
12. Huang, E. J. *et al.* Extensive FUS-Immunoreactive Pathology in Juvenile Amyotrophic Lateral Sclerosis with Basophilic Inclusions. *Brain Pathology* **20**, 1069–1076 (2010).
13. Rademakers, R. *et al.* Fus gene mutations in familial and sporadic amyotrophic lateral sclerosis. *Muscle Nerve* **42**, 170–176 (2010).
14. Suzuki, N. *et al.* FALS with FUS mutation in Japan, with early onset, rapid progress and basophilic inclusion. *J Hum Genet* **55**, 252–254 (2010).
15. Waibel, S., Neumann, M., Rabe, M., Meyer, T. & Ludolph, A. C. Novel missense and truncating mutations in FUS/TLS in familial ALS. *Neurology* **75**, 815–817 (2010).
16. Yan, J. *et al.* Frameshift and novel mutations in FUS in familial amyotrophic lateral sclerosis and ALS/dementia. *Neurology* **75**, 807–14 (2010).
17. Syriani, E., Morales, M. & Gamez, J. FUS/TLS gene mutations are the second most frequent cause of familial ALS in the Spanish population. *Amyotroph Lateral Scler* **12**, 118–23 (2011).
18. Tsai, C.-P. *et al.* FUS, TARDBP, and SOD1 mutations in a Taiwanese cohort with familial ALS. *Neurobiol Aging* **32**, 553.e13–21 (2011).
19. Belzil, V. v *et al.* Novel FUS deletion in a patient with juvenile amyotrophic lateral sclerosis. *Arch Neurol* **69**, 653–6 (2012).
20. Conte, A. *et al.* P525L FUS mutation is consistently associated with a severe form of juvenile Amyotrophic Lateral Sclerosis. *Neuromuscular Disorders* **22**, 73–75 (2012).
21. Hara, M. *et al.* Lower motor neuron disease caused by a novel FUS/TLS gene frameshift mutation. *J Neurol* **259**, 2237–9 (2012).

22. Mochizuki, Y. *et al.* Familial ALS with FUS P525L mutation: two Japanese sisters with multiple systems involvement. *J Neurol Sci* **323**, 85–92 (2012).
23. Nagayama, S. *et al.* Novel FUS mutation in patients with sporadic amyotrophic lateral sclerosis and corticobasal degeneration. *Journal of Clinical Neuroscience* **19**, 1738–1739 (2012).
24. Sproviero, W. *et al.* FUS mutations in sporadic amyotrophic lateral sclerosis: clinical and genetic analysis. *Neurobiol Aging* **33**, 837.e1–5 (2012).
25. Suzuki, N. *et al.* FUS/TLS-immunoreactive neuronal and glial cell inclusions increase with disease duration in familial amyotrophic lateral sclerosis with an R521C FUS/TLS mutation. *J Neuropathol Exp Neurol* **71**, 779–88 (2012).
26. van Blitterswijk, M. *et al.* Genetic Overlap between Apparently Sporadic Motor Neuron Diseases. *PLoS One* **7**, e48983 (2012).
27. van Blitterswijk, M. *et al.* Evidence for an oligogenic basis of amyotrophic lateral sclerosis. *Hum Mol Genet* **21**, 3776–3784 (2012).
28. Zou, Z.-Y. *et al.* Screening of the FUS gene in familial and sporadic amyotrophic lateral sclerosis patients of Chinese origin. *Eur J Neurol* **19**, 977–83 (2012).
29. Bertolin, C. *et al.* Improving the knowledge of amyotrophic lateral sclerosis genetics: Novel SOD1 and FUS variants. *Neurobiol Aging* **35**, 1212.e7-1212.e10 (2014).
30. Kent, L. *et al.* Autosomal dominant inheritance of rapidly progressive amyotrophic lateral sclerosis due to a truncation mutation in the fused in sarcoma (FUS) gene. *Amyotroph Lateral Scler Frontotemporal Degener* **15**, 557–562 (2014).
31. Mochizuki, Y. *et al.* An autopsy case of familial amyotrophic lateral sclerosis with FUS R521G mutation. *Amyotroph Lateral Scler Frontotemporal Degener* **15**, 305–8 (2014).
32. Mochizuki, Y. *et al.* A Japanese patient with familial ALS and a p.K510M mutation in the gene for FUS (FUS) resulting in the totally locked-in state. *Neuropathology* **34**, 504–9 (2014).
33. Onohara, A. *et al.* Japanese amyotrophic lateral sclerosis patient with learning disabilities with a deletion mutation in the C-terminal of the FUS/TLS gene. *Neurol Clin Neurosci* **3**, 192–193 (2015).
34. Kuang, L. *et al.* Clinical and experimental studies of a novel P525R FUS mutation in amyotrophic lateral sclerosis. *Neurol Genet* **3**, e172 (2017).
35. Özoğuz, A. *et al.* The distinct genetic pattern of ALS in Turkey and novel mutations. *Neurobiol Aging* **36**, 1764.e9-1764.e18 (2015).
36. Vance, C. *et al.* Mutations in FUS, an RNA processing protein, cause familial amyotrophic lateral sclerosis type 6. *Science* **323**, 1208–1211 (2009).
37. Murayama, S. [Clinical and pathological characteristics of FUS/TLS-associated amyotrophic lateral sclerosis (ALS)]. *Rinsho Shinkeigaku* **50**, 948–50 (2010).
38. Yamamoto-Watanabe, Y. *et al.* A Japanese ALS6 family with mutation R521C in the FUS/TLS gene: A clinical, pathological and genetic report. *J Neurol Sci* **296**, 59–63 (2010).
39. Kobayashi, Z. *et al.* Occurrence of basophilic inclusions and FUS-immunoreactive neuronal and glial inclusions in a case of familial amyotrophic lateral sclerosis. *J Neurol Sci* **293**, 6–11 (2010).
40. Chiò, A. *et al.* A de novo missense mutation of the FUS gene in a 'true' sporadic ALS case. *Neurobiol Aging* **32**, 553.e23–6 (2011).
41. Lai, S.-L. *et al.* FUS mutations in sporadic amyotrophic lateral sclerosis. *Neurobiol Aging* **32**, 550.e1–4 (2011).

42. Mackenzie, I. R. A. *et al.* Pathological heterogeneity in amyotrophic lateral sclerosis with FUS mutations: two distinct patterns correlating with disease severity and mutation. *Acta Neuropathol* **122**, 87–98 (2011).
43. Robertson, J. *et al.* A novel double mutation in FUS gene causing sporadic ALS. *Neurobiol Aging* **32**, 553.e27–30 (2011).
44. Kono, S. *et al.* Combined FDG and raclopride PET study in a case of ALS with the R521C FUS gene mutation. *J Neurol* **259**, 367–9 (2012).
45. Yamashita, S. *et al.* Sporadic juvenile amyotrophic lateral sclerosis caused by mutant FUS/TLS: possible association of mental retardation with this mutation. *J Neurol* **259**, 1039–1044 (2012).
46. Waibel, S. *et al.* Truncating mutations in FUS/TLS give rise to a more aggressive ALS-phenotype than missense mutations: a clinico-genetic study in Germany. *Eur J Neurol* **20**, 540–546 (2013).
47. Calvo, A. *et al.* A de novo nonsense mutation of the FUS gene in an apparently familial amyotrophic lateral sclerosis case. *Neurobiol Aging* **35**, 1513.e7-1513.e11 (2014).
48. Tarlarini, C. *et al.* Novel FUS mutations identified through molecular screening in a large cohort of familial and sporadic amyotrophic lateral sclerosis. *Eur J Neurol* **22**, 1474–1481 (2015).
49. Agarwal, S. *et al.* Utility of whole exome sequencing in evaluation of juvenile motor neuron disease. *Muscle Nerve* **53**, 648–652 (2016).
50. Aizawa, H. *et al.* Deficient RNA-editing enzyme ADAR2 in an amyotrophic lateral sclerosis patient with a FUSP525L mutation. *Journal of Clinical Neuroscience* **32**, 128–129 (2016).
51. Hirayanagi, K., Sato, M., Furuta, N., Makioka, K. & Ikeda, Y. Juvenile-onset Sporadic Amyotrophic Lateral Sclerosis with a Frameshift *FUS* Gene Mutation Presenting Unique Neuroradiological Findings and Cognitive Impairment. *Internal Medicine* **55**, 689–693 (2016).
52. Zou, Z.-Y., Liu, M.-S., Li, X.-G. & Cui, L.-Y. The distinctive genetic architecture of ALS in mainland China: Table 1. *J Neurol Neurosurg Psychiatry* **87**, 906–907 (2016).
53. Zou, Z.-Y., Liu, M.-S., Li, X.-G. & Cui, L.-Y. Mutations in FUS are the most frequent genetic cause in juvenile sporadic ALS patients of Chinese origin. *Amyotroph Lateral Scler Frontotemporal Degener* **17**, 249–52 (2016).
54. Gromicho, M., Oliveira Santos, M., Pinto, A., Pronto-Laborinho, A. & de Carvalho, M. Young-onset rapidly progressive ALS associated with heterozygous FUS mutation. *Amyotroph Lateral Scler Frontotemporal Degener* **18**, 451–453 (2017).
55. Belzil, V. v. *et al.* Mutations in FUS cause FALS and SALS in French and French Canadian populations. *Neurology* **73**, 1176–1179 (2009).
56. Ticozzi, N. *et al.* Analysis of FUS gene mutation in familial amyotrophic lateral sclerosis within an Italian cohort. *Neurology* **73**, 1180–1185 (2009).
57. Tateishi, T. *et al.* Multiple system degeneration with basophilic inclusions in Japanese ALS patients with FUS mutation. *Acta Neuropathol* **119**, 355–64 (2010).
58. Millecamps, S. *et al.* SOD1, ANG, VAPB, TARDBP, and FUS mutations in familial amyotrophic lateral sclerosis: genotype-phenotype correlations. *J Med Genet* **47**, 554–560 (2010).
59. Drepper, C., Herrmann, T., Wessig, C., Beck, M. & Sendtner, M. C-terminal FUS/TLS mutations in familial and sporadic ALS in Germany. *Neurobiol Aging* **32**, 548.e1–4 (2011).

60. Kwon, M. J. *et al.* Screening of the SOD1, FUS, TARDBP, ANG, and OPTN mutations in Korean patients with familial and sporadic ALS. *Neurobiol Aging* **33**, 1017.e17-1017.e23 (2012).
61. Zou, Z. Y. *et al.* De novo FUS gene mutations are associated with juvenile-onset sporadic amyotrophic lateral sclerosis in China. *Neurobiol Aging* **34**, 1312.e1-1312.e8 (2013).
62. Belzil, V. v *et al.* Identification of a FUS splicing mutation in a large family with amyotrophic lateral sclerosis. *J Hum Genet* **56**, 247–249 (2011).
63. Belzil, V. v. *et al.* Identification of novel FUS mutations in sporadic cases of amyotrophic lateral sclerosis. *Amyotrophic Lateral Sclerosis* **12**, 113–117 (2011).
64. van Langenhove, T. *et al.* Genetic contribution of FUS to frontotemporal lobar degeneration. *Neurology* **74**, 366–71 (2010).
65. Kim, Y. E. *et al.* De novo FUS mutations in 2 Korean patients with sporadic amyotrophic lateral sclerosis. *Neurobiol Aging* **36**, 1604.e17-1604.e19 (2015).
66. Hübers, A. *et al.* De novo FUS mutations are the most frequent genetic cause in early-onset German ALS patients. *Neurobiol Aging* **36**, 3117.e1-3117.e6 (2015).
67. Ito, H. *et al.* Optineurin is co-localized with FUS in basophilic inclusions of ALS with FUS mutation and in basophilic inclusion body disease. *Acta Neuropathol* **121**, 555–557 (2011).
68. Fecto, F. & Siddique, T. Making Connections: Pathology and Genetics Link Amyotrophic Lateral Sclerosis with Frontotemporal Lobe Dementia. *Journal of Molecular Neuroscience* **45**, 663–675 (2011).
69. King, A. *et al.* ALS-FUS pathology revisited: singleton FUS mutations and an unusual case with both a FUS and TARDBP mutation. *Acta Neuropathol Commun* **3**, 62 (2015).
70. Leblond, C. S. *et al.* De novo FUS P525L mutation in Juvenile amyotrophic lateral sclerosis with dysphonia and diplopia. *Neurol Genet* **2**, e63 (2016).
71. Sabatelli, M. *et al.* Mutations in the 3' untranslated region of FUS causing FUS overexpression are associated with amyotrophic lateral sclerosis. *Hum Mol Genet* **22**, 4748–55 (2013).
72. Couthouis, J., Raphael, A. R., Daneshjou, R. & Gitler, A. D. Targeted Exon Capture and Sequencing in Sporadic Amyotrophic Lateral Sclerosis. *PLoS Genet* **10**, e1004704 (2014).
73. Cady, J. *et al.* Amyotrophic lateral sclerosis onset is influenced by the burden of rare variants in known amyotrophic lateral sclerosis genes. *Ann Neurol* **77**, 100–113 (2015).
74. Dekker, A. M. *et al.* Large-scale screening in sporadic amyotrophic lateral sclerosis identifies genetic modifiers in C9orf72 repeat carriers. *Neurobiol Aging* **39**, 220.e9-220.e15 (2016).
75. Zou, Z.-Y., Liu, M.-S., Li, X.-G. & Cui, L.-Y. Mutations in SOD1 and FUS caused juvenile-onset sporadic amyotrophic lateral sclerosis with aggressive progression. *Ann Transl Med* **3**, 221 (2015).
76. Brown, J. A. *et al.* SOD1, ANG, TARDBP and FUS mutations in amyotrophic lateral sclerosis: A United States clinical testing lab experience. *Amyotrophic Lateral Sclerosis* **13**, 217–222 (2012).
77. Rutherford, N. J. *et al.* Pathogenicity of exonic indels in fused in sarcoma in amyotrophic lateral sclerosis. *Neurobiol Aging* **33**, 424.e23-424.e24 (2012).
78. Akiyama, T. *et al.* Genotype-phenotype relationships in familial amyotrophic lateral sclerosis with FUS/TLS mutations in Japan. *Muscle Nerve* **54**, 398–404 (2016).

79. Hou, L. *et al.* Screening of SOD1, FUS and TARDBP genes in patients with amyotrophic lateral sclerosis in central-southern China. *Sci Rep* **6**, 32478 (2016).
80. Kim, H.-J. *et al.* Identification of mutations in Korean patients with amyotrophic lateral sclerosis using multigene panel testing. *Neurobiol Aging* **37**, 209.e9-209.e16 (2016).
81. Corcia, P. *et al.* A novel mutation of the C-terminal amino acid of FUS (Y526C) strengthens FUS gene as the most frequent genetic factor in aggressive juvenile ALS. *Amyotroph Lateral Scler Frontotemporal Degener* **18**, 298–301 (2017).
82. Liu, Z.-J. *et al.* The investigation of genetic and clinical features in Chinese patients with juvenile amyotrophic lateral sclerosis. *Clin Genet* **92**, 267–273 (2017).
83. Müller, K. *et al.* Comprehensive analysis of the mutation spectrum in 301 German ALS families. *J Neurol Neurosurg Psychiatry* **89**, 817–827 (2018).
84. Hikiami, R. *et al.* Amyotrophic Lateral Sclerosis after Receiving the Human Papilloma Virus Vaccine: A Case Report of a 15-year-old Girl. *Intern Med* **57**, 1917–1919 (2018).
85. Yu, X. *et al.* Clinical and genetic features of patients with juvenile amyotrophic lateral sclerosis with fused in Sarcoma (FUS) mutation. *Medical Science Monitor* **24**, 8750–8757 (2018).
86. Borrego-Écija, S. *et al.* Does ALS-FUS without FUS mutation represent ALS-FET? Report of three cases. *Neuropathol Appl Neurobiol* **45**, 421–426 (2019).
87. Fujita, Y., Fujita, S., Takatama, M., Ikeda, M. & Okamoto, K. Numerous FUS-positive inclusions in an elderly woman with motor neuron disease. *Neuropathology* **31**, 170–176 (2011).
88. Takeuchi, R. *et al.* Transportin 1 accumulates in FUS inclusions in adult-onset ALS without FUS mutation. *Neuropathol Appl Neurobiol* **39**, 580–4 (2013).
89. Matsuoka, T. *et al.* An autopsied case of sporadic adult-onset amyotrophic lateral sclerosis with FUS-positive basophilic inclusions. *Neuropathology* **31**, 71–6 (2011).
90. Lattante, S. *et al.* Contribution of major amyotrophic lateral sclerosis genes to the etiology of sporadic disease. *Neurology* **79**, 66–72 (2012).
91. Dodd, K. C., Power, R., Ealing, J. & Hamdalla, H. FUS-ALS presenting with myoclonic jerks in a 17-year-old man. *Amyotroph Lateral Scler Frontotemporal Degener* **20**, 278–280 (2019).
92. Eura, N. *et al.* A juvenile sporadic amyotrophic lateral sclerosis case with P525L mutation in the FUS gene: A rare co-occurrence of autism spectrum disorder and tremor. *J Neurol Sci* **398**, 67–68 (2019).
93. Larner, A. J. & Fratalia, L. Focal limb weakness (monoparesis): when family history holds the key to diagnosis. *Br J Hosp Med (Lond)* **80**, 110–111 (2019).
94. Wharton, S. B. *et al.* Combined fused in sarcoma-positive (FUS+) basophilic inclusion body disease and atypical tauopathy presenting with an amyotrophic lateral sclerosis/motor neurone disease (ALS/MND)-plus phenotype. *Neuropathol Appl Neurobiol* **45**, 586–596 (2019).
95. Chen, L., Li, J., Lu, H. & Liu, Y. A de novo c.1509dupA:p.R503fs mutation of FUS: report of a girl with sporadic juvenile amyotrophic lateral sclerosis. *Amyotroph Lateral Scler Frontotemporal Degener* **21**, 635–637 (2020).
96. Flies, C. M. & Veldink, J. H. Chorea is a pleiotropic clinical feature of mutated fused-in-sarcoma in amyotrophic lateral sclerosis. *Amyotroph Lateral Scler Frontotemporal Degener* **21**, 309–311 (2020).

97. Picher-Martel, V., Brunet, F., Dupré, N. & Chrestian, N. The Occurrence of FUS Mutations in Pediatric Amyotrophic Lateral Sclerosis: A Case Report and Review of the Literature. *J Child Neurol* **35**, 883073820915099 (2020).
98. Wongworawat, Y. C. *et al.* Aggressive FUS-Mutant Motor Neuron Disease Without Profound Spinal Cord Pathology. *J Neuropathol Exp Neurol* **79**, 365–369 (2020).
99. Bodur, M., Toker, R. T., Başak, A. N. & Okan, M. S. A rare case of juvenile amyotrophic lateral sclerosis. *Turk J Pediatr* **63**, 495–499 (2021).
100. Lanteri, P. *et al.* The heterozygous deletion c.1509_1510delAG in exon 14 of FUS causes an aggressive childhood-onset ALS with cognitive impairment. *Neurobiol Aging* **103**, 130.e1-130.e7 (2021).
101. Moglia, C., Calvo, A., Brunetti, M., Chiò, A. & Grassano, M. Broadening the clinical spectrum of FUS mutations: a case with monomelic amyotrophy with a late progression to amyotrophic lateral sclerosis. *Neurol Sci* **42**, 1207–1209 (2021).
102. Murakami, A. *et al.* An autopsy case report of neuronal intermediate filament inclusion disease presenting with predominantly upper motor neuron features. *Neuropathology* **41**, 357–365 (2021).
103. Tanemoto, M. *et al.* Sporadic Amyotrophic Lateral Sclerosis Due to a FUS P525L Mutation with Asymmetric Muscle Weakness and Anti-ganglioside Antibodies. *Intern Med* **60**, 1949–1953 (2021).
104. Bazán-Rodríguez, L. *et al.* FUS as a cause of familial Amyotrophic lateral sclerosis, a case report in a pregnant patient. *Neurocase* **28**, 323–330 (2022).
105. Canosa, A. *et al.* A novel splice site FUS mutation in a familial ALS case: effects on protein expression. *Amyotroph Lateral Scler Frontotemporal Degener* **23**, 128–136 (2022).
106. Fiondella, L. *et al.* Co-Occurrence of Multiple Sclerosis and Amyotrophic Lateral Sclerosis in an FUS-Mutated Patient: A Case Report. *Brain Sci* **12**, 531 (2022).
107. Martinelli, I. *et al.* G507D mutation in FUS gene causes familial amyotrophic lateral sclerosis with a specific genotype-phenotype correlation. *Neurobiol Aging* **118**, 124–128 (2022).
108. Lu, T., Yang, J., Luo, L. & Wei, D. FUS mutations in Asian amyotrophic lateral sclerosis patients: a case report and literature review of genotype-phenotype correlations. *Amyotroph Lateral Scler Frontotemporal Degener* **23**, 580–584 (2022).
109. Wu, Y., Li, C., Yang, T., Lin, J. & Shang, H. A case of juvenile-onset amyotrophic lateral sclerosis with a de novo frameshift FUS gene mutation presenting with bilateral abducens palsy. *Amyotroph Lateral Scler Frontotemporal Degener* **23**, 313–314 (2022).
110. Zhou, B. *et al.* FUS P525L mutation causing amyotrophic lateral sclerosis and movement disorders. *Brain Behav* **10**, e01625 (2020).
111. Merner, N. D. *et al.* Exome Sequencing Identifies FUS Mutations as a Cause of Essential Tremor. *The American Journal of Human Genetics* **91**, 313–319 (2012).
112. Zou, Z.-Y. *et al.* Novel FUS mutation Y526F causing rapidly progressive familial amyotrophic lateral sclerosis. *Amyotroph Lateral Scler Frontotemporal Degener* **22**, 73–79 (2021).
113. Naumann, M. *et al.* Phenotypes and malignancy risk of different FUS mutations in genetic amyotrophic lateral sclerosis. *Ann Clin Transl Neurol* **6**, 2384–2394 (2019).
114. Brenner, D. *et al.* FUS mutations dominate TBK1 mutations in FUS/TBK1 double-mutant ALS/FTD pedigrees. *Neurogenetics* **23**, 59–65 (2022).

115. Lattante, S. *et al.* Coexistence of variants in TBK1 and in other ALS-related genes elucidates an oligogenic model of pathogenesis in sporadic ALS. *Neurobiol Aging* **84**, 239.e9-239.e14 (2019).
116. Jutzi, D. *et al.* Aberrant interaction of FUS with the U1 snRNA provides a molecular mechanism of FUS induced amyotrophic lateral sclerosis. *Nat Commun* **11**, 6341 (2020).
117. Reber, S. *et al.* CRISPR-Trap: a clean approach for the generation of gene knockouts and gene replacements in human cells. *Mol Biol Cell* **29**, 75–83 (2018).
